# A peptide-signal amplification strategy for the detection and validation of neoepitope presentation on cancer biopsies

**DOI:** 10.1101/2020.06.12.145276

**Authors:** Sri H. Ramarathinam, Pouya Faridi, Angela Peng, Pacman Szeto, Nicholas C. Wong, Andreas Behren, Mark Shackleton, Anthony W. Purcell

## Abstract

Targeting the right cancer-specific peptides presented by Human Leukocyte antigen (HLA) class I and II molecules on the tumor cell surface is a crucial step in cancer immunotherapy. Numerous approaches have been proposed to predict the presentation of potential neoepitopes that may be targeted through immune-based therapies. Often founded on patient specific somatic mutations, the routine validation of their actual appearance on the tumor cell surface is a significant barrier to realising personalized cancer immunotherapy. This can be attributed to the lack of robust and adaptable assays for antigen presentation that offer the required sensitivity to deal with the limited amounts of patient tumor tissue available. Rather than personalize individual assays we propose the use mass spectrometry to identify tumor neoepitopes from HLA-bound peptides directly isolated form the surface of tumor biopsies. We have developed a microscale HLA-peptide complex immunoprecipitation protocol combined with tandem mass tagging (TMT) to directly sequence HLA-bound peptides using mass spectrometry. Using this strategy, we identified HLA-bound peptides from as few as ~1000 cultured cells and from a small piece (~1 mg) of whole melanoma tumour tissue, encompassing epitopes derived from Melanoma-associated antigens and potential neoantigens.

---

HLA-bound peptides, collectively termed the ‘immunopeptidome’, can be isolated from a variety of cells and tissues using immunoaffinity precipitation of complexes with specific antibodies [1]. However, in the current approach the depth of coverage obtained is highly dependent on the amount of starting material. Tandem mass tag (TMT) labelling is an established tool in proteomics for barcoding peptides. It consists of an array of isobaric tags that are used to label individual samples [2]. Our approach takes advantage of the additive signal of common isobaric-tagged peptides within multiple samples and the ability to deconvolute sample-specific information during fragmentation of the peptides acquired in tandem mass spectra (MS2 or MS3 modalities) [3]. Several approaches to low-cell number proteomics have been described and, recently, for instance Slavlov and colleagues used TMT labelling to measure proteins at the single cell level (Scope-MS) [4–6]. This remarkable sensitivity was obtained using the additive signal from a proteolytically digested carrier proteome derived from several hundred cells. This provided sufficient ions for identification of peptides with peptides isolated from a single cell. Here the reporter ion intensities from the barcodes (TMT tags) provided quantitative information on the levels of peptides derived from single cells.

Applying peptide-barcoding to HLA-bound peptides is challenging, and typically requires a large amount of starting material, with a recently published study using either 1e8 cells or 1-2 g of mouse tissue [7], This is because HLA proteins form a small subsection of the proteome, and the complexes of peptide antigen and HLA need to be immunoprecipitated before isolating the diverse peptide cargo for subsequent analysis by mass spectrometry [1]. Moreover, unlike analyzing global proteome levels where the choice of carrier peptidome is relatively straightforward due to the presence of shared proteins, HLA-bound peptides differ according to the immunogenetics of the individual and are derived endogenously from a diverse source of parental antigens, presenting additional challenges in choosing the right (HLA-matched) carrier peptidome.

We propose an extension of the carrier-based peptide barcoding technique with key changes to sample processing, labelling and analysis strategies to confidently identify HLA-peptides from scarce samples. To facilitate identification of HLA-peptides for potential clinical translation, we refined both the peptide barcoding and HLA-peptide elution strategies to generate a robust and sensitive protocol with a 2- to 3-day turnaround. To evaluate the level of sensitivity for this approach, immunopeptidomes isolated from 5×10^6^ (as a carrier) was added to a titration of samples of HLA-bound peptides immunoprecipitated from 5×10^6^ to 1000 IHW9033 B-Lymphoblastoid cells were isolated and processed. The cell pellets were individually lysed, followed by micro-scale immunoaffinity purification of HLA molecules using antibodies specific for HLA-class I (w6/32 pan human class I specificity) and the class II molecule HLA-DR (LB3.1 with pan HLA-DR specificity). The modified microscale HLA-peptide isolation method used microscale columns that were centrifuged to achieve rapid sample loading, washing and elution of the HLA-peptide complexes. Peptide ligands from each sample were then isolated by ultrafiltration (5KDa MWCO filters) and adjusted to pH 8 prior to being labelled using individual TMT labels (Fig 1). The samples were then combined, concentrated, desalted and analyzed on a Fusion Tribrid mass spectrometer using synchronous precursor selection combined with MS3 (SPS-MS3) to reduce coisolation and compression of TMT tag ratios [3].

**Figure 1:**
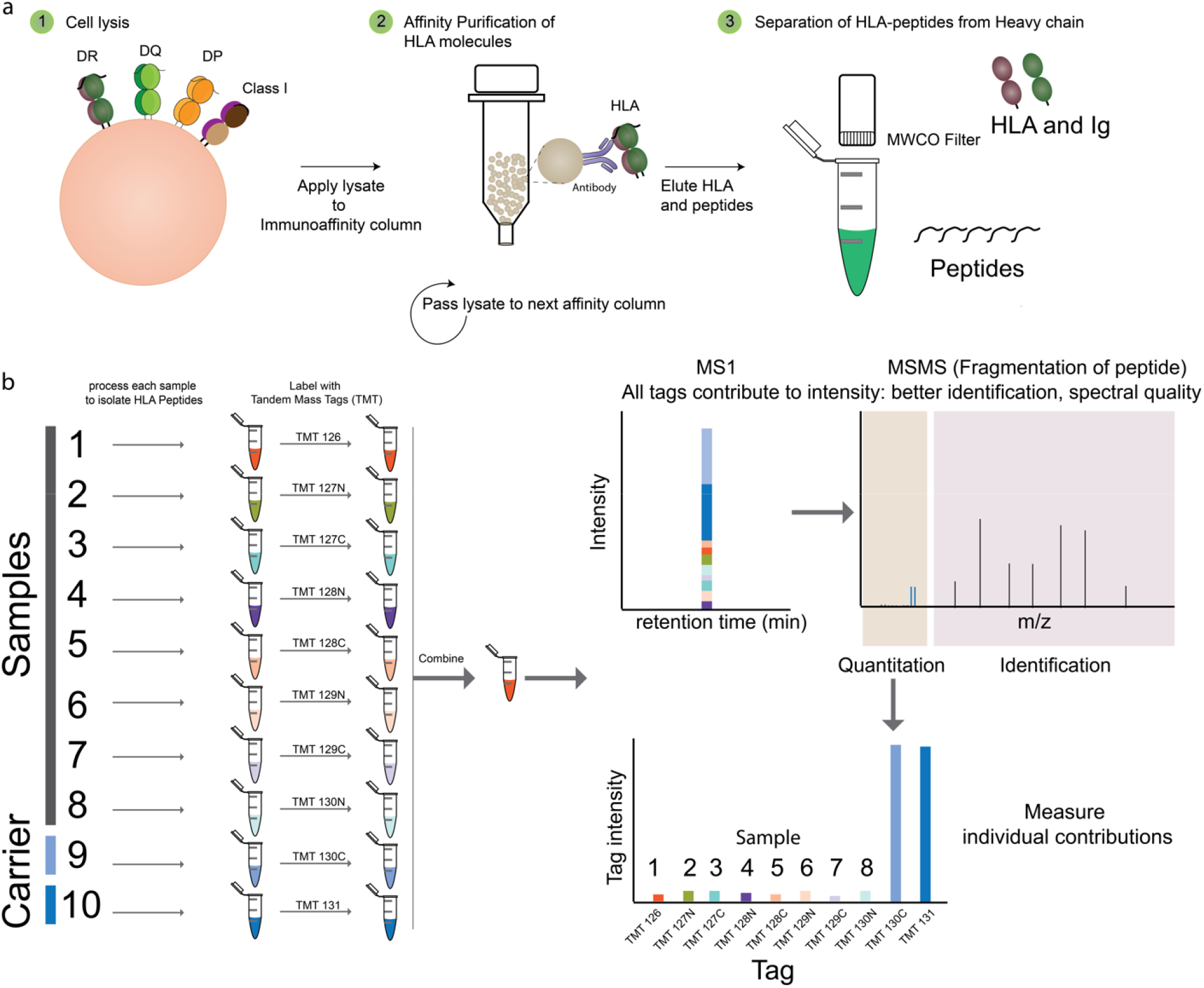
Identification of HLA peptides from scarce samples by peptide barcoding. A) The samples are lysed and the HLA molecules affinity purified using antibodies coupled to either magnetic beads or agarose beads coupled with protein A. The Peptides are then separated from HLA molecules and antibodies using molecular weight cut off filters. B) The eluted HLA peptides from the scarce samples and carrier samples are neutralized, tagged with a specific TMT channel peptide of interest prior to analysis by high resolution mass spectrometry using synchronous precursor scanning (SPS) -MS3 method. The carrier peptides provide MSMS for identification (MS2) and the contribution of each channel of TMT at the MS3 level would correspond to amount of peptide present in each sample.

As anticipated, by decreasing cellular input the signal intensity of TMT rapidly dropped. However due to the abundance of carrier peptides, reliable signals for many peptides were still evident even with as few as 1000 cells (Fig 2A). With this level of cellular input, 688 HLA-bound peptides from HLA class I and HLA-DR were identified at a false discovery rate (FDR) of 1%. The carrier peptidome sample (derived from 5e6 cells) consisted of 861 peptides (Supp Table 1) of which 80% were identified from 1000 cells. The length distribution of identified peptides was typical of HLA peptides with the majority of peptides within 8-12 mers for class I and 14-18 mers for class II. The peptide sequences identified by this technique contain the expected binding motif of HLA molecules expressed in these cells (Fig 2B, C) strengthening the confidence in the identified peptides.

**Figure 2:**
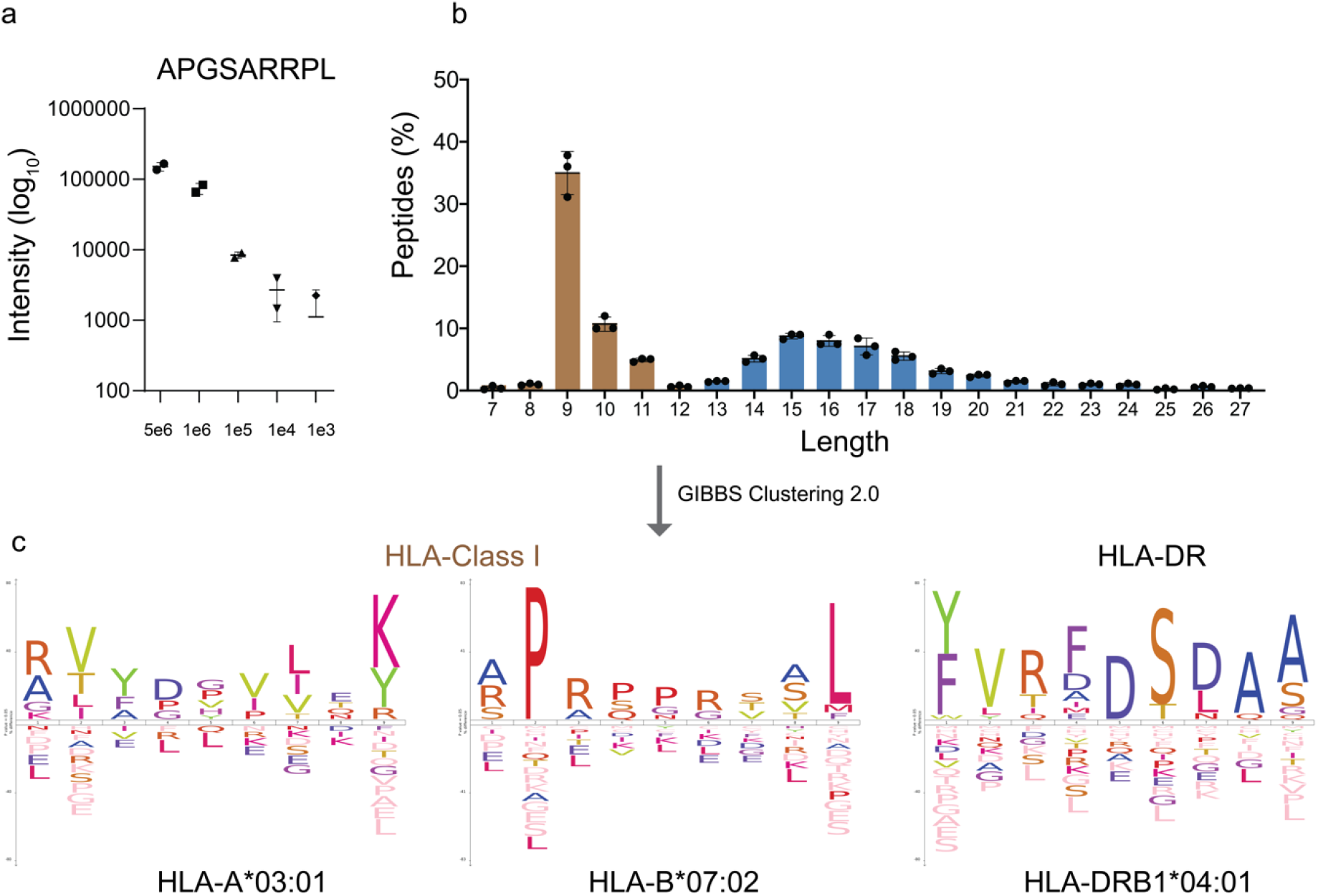
Identification of HLA-peptides from 1000 cells. A) The BLCLs (9033 cell line) were titrated from 5e6 to 1000 cells in duplicate and labelled using TMT10plex reagents. The intensity of the TMT tags is shown for a HLA class I peptide. B) The length distribution of the HLA class I (brown) and class II (blue) fall within typical ranges expected. Error bars represent mean+/-SD C) Clustering of the peptides can distinguish the HLA class I alleles expressed by this cell line (HLA-A*03:01, −B*07:02 and HLA-DRB1*04:01) that were isolated using class I -and DR-specific antibodies w6/32 and L3.1 and match the typical binding motif of these alleles.

To apply this strategy to a context of potential clinical relevance, a biopsy from a HLA-A2^+^ melanoma was obtained along with a corresponding patient-derived xenograft (PDX) from the same individual. Two recent studies have shown the importance of PDX-based models in studying immunopeptidomics establishing their role as an appropriate carrier for studying small biopsies[8, 9]. Melanoma PDX can be readily developed and expanded in NOD.SCID (NSG) mice [10] to provide an abundant source of peptide ligands and act as a carrier immunopeptidome for personalized neoepitope identification. To identify HLA class I peptides from 19 mg or 1 mg of melanoma biopsy, differing amounts (400 mg, 150 mg and 50 mg) of PDX tissue were used as carrier tags in duplicate (See Tag information in Supplementary data file for experimental setup). The samples were then processed and labelled as described above. The tagged carrier peptides from PDX provide necessary intensity of ions to aid in identifying the peptides from the sample-limited biopsy when combined and analyzed by mass spectrometry. Based on this analysis, 1257 peptides presented by HLA class I molecules were identified in the patient melanoma biopsy, and 1944 peptides were identified in the larger mass (400 mg) of PDX tumor tissue, as determined using conventional LC–MS/MS of TMT-labelled peptides (Supp Table 2). Most peptides were 8-12 mers (Fig 3b), and had the expected binding motif for the HLA molecules of the patient. Analysis of the overall intensities of peptides represented by each label tag showed the expected distribution of TMT intensities with the PDX tumor yielding the most peptide (Fig 3c). Of note, some epitopes were disproportionally highly abundant in the patient biopsy compared to the PDX tumor (see heatmap of epitopes vs other peptides Fig 3c). This suggests that there may be differences in epitope densities in tumors in patients compared to the PDX tumors. Moreover, while most peptides (88%) had HLA-binding rank (NetMHCpan4.0) of ≤2, several ligands eluted directly from the patient biopsy were within the percentile rank range 2-5 that would have been missed if prediction software [11] were used to shortlist candidate epitopes.

**Figure 3:**
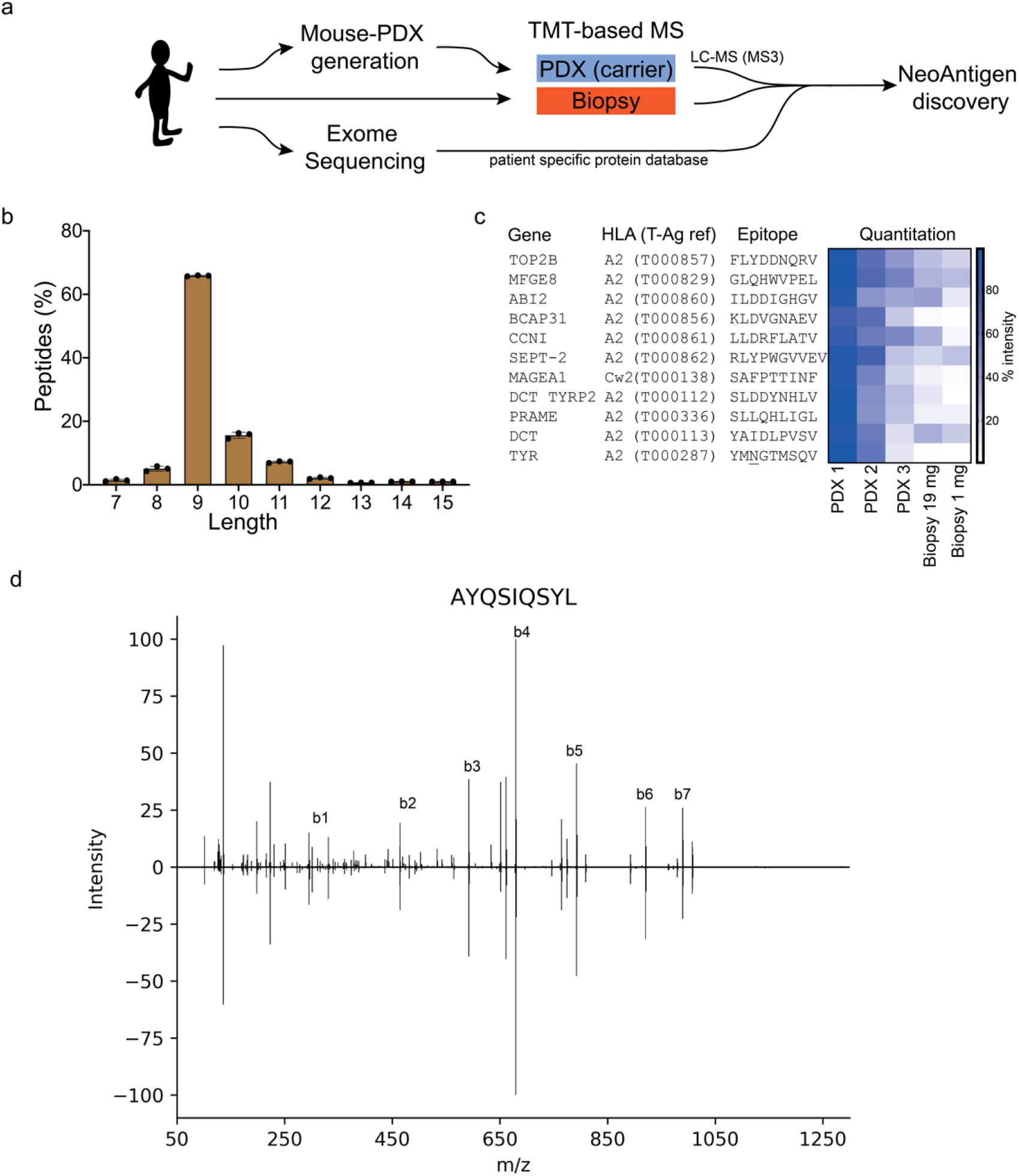
Identification of neoepitopes from patient biopsies. A) HLA peptides were identified from either 19 mg or 1 mg of Patient biopsy (HLA A2) by using a patient derived xenograft (PDX) as carrier. The Exome sequencing allowed generation of a patient-specific database to search for mutated HLA-peptides. B) The HLA class I peptides isolated followed typical length range (8-12 mers). Error bars represent mean+/-SD. C) Several known HLA-A2 epitopes from cancer associated proteins were identified in the biopsy. D) Use of patient-specific protein database helped identify a neo-antigen arising from a N to S mutation (N401S) in ER Degradation Enhancing Alpha-Mannosidase Like Protein EDEM1) which was subsequently validated using synthetic peptide (Pearson correlation r 0.9961 [CI 0.9780 to 0.9993] for b and y ions).

Analysis of peptides identified in patient biopsies revealed expression of 11 known HLA-A2 restricted tumor epitopes listed in the T-Antigen database (Supp Table 3), 102 peptides from 61 proteins sources in the T-antigen database (Supp Table 4), and 11 peptides from proteins in the Cancer Testis Antigen database (CTA; Supp Table 5). Seventeen peptides from proteins associated with melanoma and other cancers, including Melanoma-associated antigens (MAGE) 1-3, L-dopachrome tautomerase (TYRP2), Preferentially expressed antigen of Melanoma (PRAME), Protein S100-A1/B (S10A1, S100B), Integrin alpha-5 (ITA5), and beta-catenin (CTNNB1) (Supp Table 6-7), were identified in the patient biopsy. Additionally, 68 peptides from 95 proteins with keywords associated with melanoma in the uniprot database (Supp Table 7) were identified in the patient biopsy.

To evaluate our strategy for detection of neoantigens (HLA bound peptides carrying somatic mutations), the exome of the melanoma biopsy was identified using Next Generation sequencing (NGS) to generate a reference proteome specific for the tumor containing variants and mutations. This was cross-referenced against the patient biopsy HLA-ligandome data, revealing 6 epitopes that spanned mutated regions including one in EDEM1 (N401S) that was verified using a synthetic peptide (Figure 3d; Supp Table 8). Identification of neoepitopes from such high-resolution data can directly inform the generation of targets for immunotherapy and highlights the utility of our approach for directly sequencing neoepitopes from even limited amount of patient biopsies.

In summary, we describe here a novel method to isolate and identify HLA-class I- and II-bound peptides from small numbers of tumor cells or small pieces of tumour tissue. By using up to 10-15 tags (using TMT-11plex or-16plex labelling reagents) to represent 10-15 clinical samples and controls, our approach enables rapid neoepitope discovery in cancers in a manner that is translatable to contexts of routine clinical care of patients. For example, as a typical fine needle tumour aspirate, a simple and widely applied method of tumour sampling in patients, contains ~1 – 5 million cells [12], our technique would allow for rapid and sensitive identification from such samples of epitopes derived from tumor-associated antigens

Moreover, we envision this strategy will be useful for studying peptidome subsets in other clinical samples where the amount of tissue available via sampling is limiting, including biopsies taken for autoimmune diseases and infections, or to sample rare cell types in tissue/cell samples, such as different antigen-presenting cell subsets. The short turnaround time of 2-3 days for this method makes it possible to get clinically relevant information for patients rapidly and can be further reduced using automated liquid handling. The technique could also be applied to study post-transcriptional modifications of peptides [13], including splicing variants [9, 14], which cannot currently be discerned by prospective isolation. Instead of relying on antigen prediction algorithms alone, our methodology can directly isolate antigens that are already presented by HLA molecules, thereby increasing the precision of personalized approaches to cancer treatment.

## Methods

### PDX, Cell lines and Biopsy

For Patient Derived Xenograft (PDX) tumour generation, de-identified fresh patient tumor specimens were obtained through the Victorian Cancer Biobank. The use of all human specimens was performed with ethical approval from the Peter MacCallum Cancer Center Human Research Ethics Committee (HREC) and the Alfred Hospital HREC. Freshly isolated or DMSO frozen patient melanoma cells were mixed with growth factor reduced Matrigel (Corning, NY, USA) in a 1:1 ratio and injected subcutaneously into NOD.Cg-Prkdcscid Il2rgtm1Wjl/SzJ (NSG) mice. NSG mice were obtained from Jackson Laboratory and both male and female mice were used. All mouse experiments were performed under protocols approved by the Alfred Research Alliance Animal Ethics Committee. PDX tumours were resected, weighed and snap frozen in liquid nitrogen. BM14 (ECACC 88052033) human HLA-Typed Lymphoblastoid Cell Line was obtained from ECACC and maintained in RPMI (ThermoFisher) media supplemented with 10% FBS in the absence of antibiotics. Cells were counted and subjected to lysis and affinity purification on the same day.

### Affinity purification of HLA-peptide complexes

HLA-bound peptides were isolated as described previously [1] with modifications as detailed below. HLA-complexes from BLCLs (IHW9033), PDX and Biopsy material were affinity captured by immunoaffinity resin (Agarose-Protein A) containing w6/32 antibody following lysis in buffer containing 0.5% IGEPAL, 50 mM TIS pH 8.0, 150 mM NaCl, 1x Protease inhibitor cocktail). The affinity capture resin was the transferred to a mobi-spin column and washed three times with 500 uL 1x PBS by centrifugation. The HLA-peptide complexes were eluted using 10% acetic acid and heated to 70°C for 10 min. The mixture containing HLA heavy chains, antibody and peptides was subjected to a 5 KDa molecular weight cut off filter (MWCO) to separate peptides from HLA molecules and antibodies. The peptides were then neutralized using 1 M triethylammonium bicarbonate (TEAB) buffer to bring pH to 8 to enable TMT labelling. Individual samples were labelled following manufacturer’s instructions and quenched using 5% hydroxylamine. The samples were then combined and concentrated using C18 column prior to centrifugal evaporation to reduce volume and ACN concentration. The peptides from this combined sample were reconstituted in 2% ACN, 0.1% formic acid prior to analysis by mass spectrometry.

### Mass spectrometry

TMT-labelled peptides were analyzed using Orbitrap Tribrid Fusion mass spectrometer (Thermo Scientific) coupled with a RSLC nano-HPLC (Ultimate 3000, Thermo Scientific). Samples were loaded on to a 100 uM, 2 cm PepMap100 trap column in 2% ACN, 0.1% formic acid at a flow rate of 15 ul/min. Peptides were then eluted at a flow rate of 250 ul/min with starting conditions of 98% Buffer A (0.1% formic acid) and 2% Buffer B (80% ACN, 0.1% formic acid) for 2 min; Buffer B was then elevated from 2% to 7.5% B over 1 min, followed by linear gradient from 7.5% to 37.5% B over 120 min, increasing to 42.5% B over 3 min, an additional increase to 99% B at the end of gradient for 6 min followed by reduction to 2% B to allow re-equilibration. The Orbitrap Fusion instrument was operated in a data-dependent acquisition mode utilizing synchronous precursor selection (SPS) as described previously with modifications to allow efficient detection of HLA peptides: Survey full scan spectra (m/z 380-1580) were acquired in the Orbitrap at 120,000 resolution at m/z 200 after accumulation of ions to a 4e5 target value with maximum injection time of 50 ms. Dynamic exclusion was set to 15 s. Ions with 2+ to 6 + charge states were selected for msms fragmentation and to enable collection of singly charged species of interest while eliminating noise, a decision tree was included to only fragment 1+ species above 800 m/z. In both cases, msms fragments were collected in orbitrap at 60,000 resolution with first mass set to 100 m/z, target of 2e5 ions, with maximum injection time of 120 ms. To quantitate the TMT-labelled peptides confidently and to reduce interference, synchronous precursor scans (10 scans) were performed on each msms spectra to select 10 peaks for further MS3 fragmentation and analyzed in Orbitrap at 60,000 resolution.

### Data analysis

TMT-MS3 LC-MS data was analyzed using Peaks X (BSI) database search against human proteome (uniprot v03_2019) and translated exome-seq data [15] with following search parameters: parent mass error tolerance for parent and fragment mass were set to 10 ppm and 0.02 Da respectively; digestion mode was set to unspecific, with TMT-10plex as fixed modification and Oxidation(M), Acetylation (N-term and K), as variable modifications (a maximum of 3 per sequence) with False Discovery Rate (FDR) of 1%. To enable assignment of MS3 reporter ion data to respective samples, PeaksQ. quantitation was used with a tight quantification mass tolerance of 3.0 ppm utilising peptides that matched the FDR threshold of 1%. Reporter ion intensities were corrected for isotopic impurities according to lot-specific manufacturer specifications. An intensity-based threshold of 500 counts in the MS3 channel were used to assign peptides to specific tags. Peptides and proteins identified were annotated with information from T-Antigen database [16], CTA [17] and uniprot [18] to identify known antigens and epitopes.

### Exome Analysis

Exome-seq on DNA extracted from the melanoma biopsy was performed by the Peter MacCallum Cancer Centre Sequencing Core on an Illumina HiSeq according to standard protocols. Data was aligned to the human reference genome g1k_v37, using bwa v0.7.12-r1039. Resultant BAM files were then processed with GATK4 [19] and MuTect [20] to call variants. Annovar [21] was then used to annotate mutations. All non-synonymous mutations were extracted and the amino acid sequence surrounding the mutations were extracted to generate a putative reference proteome from the sample.

## Acknowledgements

The authors thank Andreas Huhmer and Daniel Lopez of ThermoScientific for helpful discussions and advice on TMT labelling of small samples.

This work was supported by the Australian National Health and Medical Research Council (NHMRC) Project (1165490). A.W.P. is supported by a NHMRC Principal Research Fellowship (1137739). We thank staff at the Monash Biomedical Proteomics Facility for technical assistance. Computational resources were supported by the R@CMon/Monash Node of the NeCTAR Research Cloud, an initiative of the Australian Government’s Super Science Scheme and the Education Investment Fund.

## References

1. Purcell, A.W., S.H. Ramarathinam, and N. Ternette, Mass spectrometry-bosed identification of MHC-bound peptides for immunopeptidomics. Nat Protoc, 2019. 14(6): p. 1687–1707.

2. Thompson, A., et al., Tandem mass tags: a novel quantification strategy for comparative analysis of complex protein mixtures by MS/MS. Anal Chem, 2003. 75(8): p. 1895–904.

3. McAlister, G.C., et al., MultiNotch MS3 enables accurate, sensitive, and multiplexed detection of differential expression across cancer cell line proteomes. Anal Chem, 2014. 86(14): p. 7150–8.

4. Zhu, Y., et al., Nanodroplet processing platform for deep and quantitative proteome profiling of 10-100 mammalian cells. Nat Commun, 2018. 9(1): p. 882.

5. Yi, L., et al., Boosting to Amplify Signal with Isobaric Labeling (BASIL) Strategy for Comprehensive Quantitative Phosphoproteomic Characterization of Small Populations of Cells. 2019.

6. Budnik, B., et al., SCoPE-MS: mass spectrometry of single mammalian cells quantifies proteome heterogeneity during cell differentiation. Genome Biol, 2018. 19(1): p. 161.

7. Murphy, J.P., et al., Multiplexed Relative Quantitation with Isobaric Tagging Mass Spectrometry Reveals Class I Major Histocompatibility Complex Ligand Dynamics in Response to Doxorubicin. Anal Chem, 2019. 91(8): p. 5106–5115.

8. Rijensky, N.M., et al., Identification of tumor antigens in the HLA peptidome of patient-derived xenograft tumors in mouse. Mol Cell Proteomics, 2020.

9. Faridi, P., et al., A subset ofHLA-l peptides are not genomically templated: Evidence for cis- and trans-spliced peptide ligands. Sci Immunol, 2018. 3(28).

10. Quintana, E., et al., Efficient tumour formation by single human melanoma cells. Nature, 2008. 456(7222): p. 593–598.

11. Peters, B., M. Nielsen, and A. Sette, T Cell Epitope Predictions. Annual Review of Immunology, 2020. 38(1): p. 123–145.

12. Rajer, M. and M. Kmet, Quantitative analysis of fine needle aspiration biopsy samples. Radiol Oncol, 2005. 39(4): p. 269–272.

13. Ramarathinam, S.H., et al., Employing proteomics in the study of antigen presentation: an update. Expert Rev Proteomics, 2018. 15(8): p. 637–645.

14. Liepe, J., et al., A large fraction of HLA class I ligands are proteasome-generated spliced peptides. 2016.

15. Tran, N.H., et al., Deep learning enables de novo peptide seguencing from data-independent-acguisition mass spectrometry. Nat Methods, 2019. 16(1): p. 63–66.

16. Olsen, L.R., et al., TANTIGEN: a comprehensive database of tumor T cell antigens. Cancer Immunol Immunother, 2017. 66(6): p. 731–735.

17. Almeida, L.G., et al., CTdatabase: a knowledge-base of high-throughput and curated data on cancer-testis antigens. Nucleic Acids Res, 2009. 37(Database issue): p. D816–9.

18. UniProt, C., UniProt: a worldwide hub of protein knowledge. Nucleic Acids Res, 2019. 47(D1): p. D506–D515.

19. McKenna, A., et al., The Genome Analysis Toolkit: A MapReduce framework for analyzing nextgeneration DNA sequencing data. Genome Research, 2010. 20(9): p. 1297–1303.

20. Cibulskis, K., et al., Sensitive detection of somatic point mutations in impure and heterogeneous cancer samples. Nature Biotechnology, 2013. 31(3): p. 213–219.

21. Wang, K., M. Li, and H. Hakonarson, ANNOVAR: functional annotation of genetic variants from high-throughput sequencing data. Nucleic Acids Research, 2010. 38(16): p. e164–e164.

